# Evaluating the efficacy of LUMPY-NIL herbal powder, dermal Spray and dermal ointment in treating Lumpy Skin Disease (LSD) in Bovines

**DOI:** 10.1101/2025.05.22.655490

**Authors:** Arun HS Kumar, Vasanthkumar Sagar, BM Ravindranath

## Abstract

**Background:** Lumpy Skin Disease (LSD) lacks targeted treatment options and is managed using antibiotics and anti-inflammatory drugs. This study evaluated the efficacy of a polyherbal combination therapy in the management of LSD in affected bovines.

**Materials and Methods:** 52 dairy cattle clinically diagnosed with LSD were enrolled in a field trial conducted under veterinary supervision. All animals received LUMPY-NIL Herbal Powder orally at 30 g twice daily, along with topical application of LUMPY-NIL Dermal Spray on closed nodules and LUMPY-NIL Ointment on open wounds. Daily monitoring included clinical signs, lesion progression, systemic symptoms, and milk yield recovery.

**Results:** Treatment duration varied from 7 to 21 days depending on severity of the disease. New lesion formation ceased within 2–4 days of treatment initiation. Systemic symptoms such as fever, lethargy, and inappetence resolved within 1–3 days. Open wounds showed progressive healing with no signs of infection, and nodules dried and sloughed off by day 15 in most animals. Milk yield showed partial recovery (20%) by day 7 and returned to baseline by day 15. The reduction in nodule size and number ranged from 81% to 94% by days 10-15. No adverse effects were observed during the treatment period.

**Conclusion:** The polyherbal treatment combining demonstrated promising therapeutic benefits in managing clinical symptoms and accelerating recovery in LSD-infected cattle.

## 1. Introduction

Lumpy skin disease (LSD) is a viral disease of cattle that is caused by the Lumpy skin disease virus (LSDV). The virus belongs to the family Poxviridae, which also includes other poxviruses that affect animals and humans. LSDV is a double-stranded DNA virus that is transmitted by blood-feeding insects, particularly certain species of mosquitoes and flies [1,2]. Clinical signs of LSD typically include fever, anorexia, and respiratory signs such as coughing and nasal discharge. The hallmark symptom of LSD is the formation of nodules or lumps on the skin, which can be accompanied by oedema (swelling) and ulceration [3,4]. The nodules can occur anywhere on the body, including the head, neck, trunk, and limbs, and can range in size from a few millimetres to several centimetres in diameter. The nodules can cause significant pain and discomfort to affected animals, and can also result in secondary infections and loss of production due to reduced mobility and feeding. These nodules can become ulcerated and may be accompanied by swelling of the lymph nodes. In severe cases, the disease can lead to death, particularly in young or immunocompromised animals. The disease can have a significant impact on animal welfare, as well as on the productivity and profitability of affected farms [5]. In addition to its impact on animal health, LSD also has significant economic consequences, particularly in affected countries where cattle are an important source of income and livelihoods [6,7]. The disease can lead to reduced milk and meat production, as well as trade restrictions on affected countries [3,8].

LSDV can also infect other ruminants, such as water buffalo, goats, and sheep [2]. The virus is relatively stable in the environment and can survive for extended periods of time in contaminated soil, water, or on fomites. The virus is primarily spread through biting insects, such as the mosquito (*Aedes aegypti, Culex quinquefasciatus*, and *Anopheles stephensi*) or the stable fly (*Stomoxys calcitrans*), as well as through contact with contaminated materials such as equipment, clothing, or feed [1,2]. The severity of the disease can vary depending on the strain of LSDV, the immune status of the animal, and other factors such as environmental conditions and concurrent infections. Mortality rates in affected herds can range from 1% to 10%, and morbidity rates can be as high as 100%, depending on the severity of the outbreak.[9,10] Diagnosis of LSD is based on clinical signs, laboratory testing, and epidemiological investigation. Laboratory diagnosis can be achieved through the isolation of the virus from skin nodules, or through detection of viral DNA or proteins using molecular methods such as polymerase chain reaction (PCR) or enzyme-linked immunosorbent assay (ELISA) [9].

LSD was first reported in Zambia in 1929,[11] and since then has been reported in several African countries, as well as in the Middle East, Europe, and Asia [12]. The disease is considered endemic in many African countries, particularly in sub-Saharan Africa [13]. However, in recent years, LSD has been spreading to new areas, including Europe, where it was first reported in Greece in 2015 and has since been reported in several other countries [4,14]. There is currently no specific treatment for LSD, and control measures rely mainly on vaccination, insect control, and quarantine of affected animals.[9] There are currently a few vaccines available for LSD, including live-attenuated vaccines, inactivated vaccines, and recombinant vaccines [15,16]. Vaccines have shown to be effective in preventing disease and reducing its spread. Studies have demonstrated that vaccination can significantly reduce the incidence and severity of LSD, reduce the number of animals affected and reduce the shedding of virus, which can help to reduce the risk of transmission to other animals.[6,15] However, vaccination coverage in many affected countries is often limited, and there are concerns about the potential for vaccine failure due to the genetic diversity of the virus.[6,15,17] The LSDV has a high level of genetic diversity, with multiple strains circulating in different regions. This genetic diversity can make it difficult to develop vaccines that are effective against all strains of the virus. In addition, the effectiveness of vaccines can be influenced by factors such as the timing and frequency of vaccination, the quality of the vaccine, and the immune status of the animals being vaccinated.[18]

There is currently no specific treatment for LSD in cattle. As the disease is caused by a virus, the treatment options are limited to supportive care, management of clinical signs, and prevention of secondary infections. Management of clinical signs in affected animals includes the use of pain relief medication to alleviate discomfort associated with skin nodules and oedema.[9,19] Antibiotics may be used to treat secondary bacterial infections that can occur in open wounds or ulcers associated with the skin nodules. Fly control measures are also important to prevent further spread of the disease and to reduce the risk of secondary infections. Several phytochemicals are reported to have antiviral activity against poxviruses and in the past have provided leads for discovery and development of effective therapeutics.[20,21] About fourteen different family of Indian plants are reported to be bioactive against poxviruses.[21-23] Evaluation of these plant extracts against specific poxviruses may lead to identification of effective therapeutics. The plant extracts are of specific interest against poxviruses as the diverse phytochemicals in the extract may effectively target the complex morphogenesis of the poxviruses. One aspect of the complex morphogenesis of the poxvirus is the prevalence of viral genome which encode for several proteins with capabilities to interact with the host processes necessary for viral replication and curtail host defence system.[22,23] Besides this, poxviruses also exhibit considerable diversity in routes through which they can enter host cells, which again merits use of plant extract for effective targeting. Although most poxviruses have similarities in their pathophysiology, a detailed understanding of LSDV specific pathophysiology is lacking, which is perhaps the reason for lack of efforts to develop effective therapeutics against it. Using a homology modelling approach, along with established molecular modelling tools and in silico pharmacology, we identified effective phytochemicals against LSDV. Based on these findings, a polyherbal formulation was developed and subsequently evaluated for its therapeutic efficacy in clinical cases of LSDV.

## 2. Materials and Methods

The 3D structure of LSDV proteins (Poly-A-polymerase catalytic subunit (Q91MX5), Envelope protein A28 homolog (Q8JTR0), GTP--RNA guanylyltransferase (Q91MT3), DNA-directed RNA polymerase (Q91MR7), host range protein 2 (Q91MU3), Envelope protein (A0A1B3B646) were downloaded as PDB files from the protein data bank (https://www.rcsb.org/), uniport database (https://www.uniprot.org/peptidesearch/) or homology modelled using the SWISS-MODEL server (https://swissmodel.expasy.org/) and were optimized for molecular docking in the Chimera software.[24-28]

The structures of selected natural compounds in *Azadirachta indica* and *Eugenia jambolana, Piper longum, Ocimum sanctum and Tinospora cordifolia* (sitosterol, betulinic acid, crategolic acid, hepatcosane, n-nonacosane, nhentriacontane, noctacosanol, ntriacontanol, ndotricontanol, mearnsetin, myricetin 3-O- (4”- acetyl-2”-O-galloyl) alpha, quercetin, myricetin, myricitrin. epoxyazadiradione, gedunin, azadirachtin, nimbin and maslinic acid) were accessed from PubChem database and processed into PDB file format and minimised for molecular docking using the Chimera software.[24,29] These natural compounds were identified from an initial inhouse hit screening scores against the LSDV targets. Molecular docking was performed to evaluate the binding efficacy of these compounds against the LSDV protein targets using AutoDock Vina (version 1.5.4) and the docked protein-ligand complex were visualised using the Chimera and PyMOL v 1.8.2.0 software[24,29-31]. AutoDock-MGLTools was employed to visualize and modify the receptor and ligand structures to PDBQT file formats. The PDBQT file formats of the ligand and receptor were used for molecular docking using the AutoDock Vina program. Ligands were docked individually to the receptor with grid coordinates and grid boxes of specific sizes for each receptor centralised in the AutoDock-MGLTools. The output file was saved in the PDBQT format and the ligand-receptor binding affinity estimated as negative Gibbs free energy (ΔG) scores (Kcal/mol), were documented based on AutoDock Vina scoring function. Post-docking analyses were visualized using PyMOL and Chimera, giving details of the sizes, locations of binding sites, hydrogen-bond interactions of the docked ligand in various confirmations.

### Simulation of dose response curves

Dose-response curves were modelled based on nonlinear regression analysis approach by specifying IC_50_ values as independent variable and response (% inhibition) as dependent variable. IC_50_ values were estimated from the binding affinity values using the following formula IC_50_ = exp(deltaG/RT) (1+ ([S]/Km)). Where deltaG = Binding affinity (Kcal/mol), RT = 298K, S = substrate concentration, K_m_=Michaelis constant. The increase in the ligand dose results in sequential changes to the response (% receptor inhibition) eventually achieving minimum or maximum response limits. As the 20 to 80% response is linear, this was modelled using four-point logistic increments of IC_50_ values (0.5x, 1x, 1.5x and 2x) and averaged. The resulting equation from this was y = 26.362x 158.85 (R^2^ = 0.9967). The one standard deviation increase and decrease of the IC_50_ values was estimated from this equation. The resulted three IC_50_ values estimated were used to calculate the IC 5, 20, 60, 80 and 100 values by employing a five-parameter logistic equation.[32] The means of the IC 5, 20, 60, 80 and 100 values obtained (in x axis) were plotted against the % inhibition response (in y axis) to obtain the simulated dose response curves.[27,33]

### Selection of medicinal plants

A panel of medicinal plants known for their antiviral, immunomodulatory, antioxidant, and wound-healing properties were selected based on existing scientific literature and affinity analysis of compounds against LSDV structural and non-structural proteins involved in viral entry (e.g., envelope proteins) and replication (e.g., DNA polymerase). Compounds demonstrating strong binding efficacy (IC_50_ range: 0.24–14.52 µM) were used as a bench mark for selecting medicinal plants for formulation development. The selected species included Cissus quadrangularis, Azadirachta indica, Glycyrrhiza glabra, Tinospora cordifolia, Ocimum sanctum, Cocos nucifera, Allium sativum, Pongamia pinnata, Piper longum, Eugenia jambolana, Curcuma longa, and Eucalyptus globulus.

### Formulation of LUMPY-NIL Herbal Powder (Oral Suspension)

To develop an oral dry suspension containing active phytochemical ingredients (APIs) capable of inhibiting LSDV replication and boosting non-specific immunity in infected cattle, the dried plant materials were powdered and subjected to hydroalcoholic extraction (70:30 ethanol:water) via Soxhlet apparatus for 6–8 hours. Extracts were filtered and concentrated under reduced pressure using a rotary evaporator. Extracts were standardized based on marker compounds (e.g., curcumin from Curcuma longa, glycyrrhizin from Glycyrrhiza glabra, eugenol from Ocimum sanctum) using HPLC analysis. Standardized extracts were blended in predetermined ratios based on IC_50_ values, synergistic effects, and literature-based therapeutic doses. Natural excipients such as Acacia gum and maltodextrin were added for flowability and palatability. Antioxidant stabilizers (e.g., ascorbic acid) were included to maintain phytochemical activity. The blended formulation was spray-dried to obtain a free-flowing powder. The powder was designed to be reconstituted in water or suitable excipient (jaggery or banana) prior to oral administration.

### Formulation of LUMPY-NIL Dermal Spray (Essential Oil-Based)

To develop a topical dermal spray composed of essential oils with antimicrobial, antiviral, and soothing properties for managing cutaneous lesions in LSD, essential oils from Eucalyptus globulus, Ocimum sanctum, Pongamia pinnata, and Cocos nucifera were obtained through steam distillation. Oils were blended using natural emulsifiers (e.g., lecithin) and solubilized in an aqueous base containing mild surfactants. Polysorbate 20 was used to enhance oil dispersion. Natural humectants (e.g., glycerin), preservatives (e.g., sodium benzoate), and soothing agents (e.g., aloe vera gel) were added. The solution was filtered through 0.22-micron filters and filled into sterilized spray bottles under aseptic conditions.

### Formulation of LUMPY-NIL Ointment (Herbal Extract-Based)

The ointment for application on cutaneous nodules and scabs caused by LSD was developed using aqueous and ethanolic extracts of Azadirachta indica, Glycyrrhiza glabra, Curcuma longa, Eugenia jambolana, and Tinospora cordifolia. A natural lipid-based ointment base was formulated using beeswax and coconut oil, to provide occlusive, healing properties. Concentrated extracts were slowly mixed into the melted base at controlled temperatures (40–50°C) to preserve phytochemical integrity. The formulation was homogenized and allowed to cool with continuous stirring to ensure uniform dispersion.

Stability testing of all three formulations was conducted over 180 days (accelerated stability study) under various storage conditions. All formulations were subjected to initial in vitro and in vivo safety screening (Acute Oral Toxicity Study of in Rats as per OECD Guideline No. 423 - Acute Toxic Class Method) for dermal tolerance, palatability (for oral powder), and systemic adverse effects. All three formulations showed acceptable safety profiles and hence were approved for field efficacy evaluation in LSDV-infected animals. All the three formulations were developed by Sparconn Life Sciences, Karnataka, India, under AYUSH license no AUS-990.

### Quality Control and Standardization Procedures

To ensure consistency, safety, and efficacy of the LUMPY-NIL Herbal Powder, Dermal Spray, and Ointment formulations, comprehensive quality control and standardization protocols were established at each stage of the development process. All medicinal plant materials were authenticated by a qualified botanist using macroscopic and microscopic characteristics. Raw plant materials were tested for heavy metals (Pb, Cd, Hg, As) using Atomic Absorption Spectroscopy (AAS), pesticide residues using GC-MS and microbial load (TVC, yeast, mould, E. coli, Salmonella) per WHO guidelines. Moisture Content was determined using a moisture analyser to prevent microbial growth and degradation. Active extracts were standardised based on phytochemical markers known to correlate with biological activity: Curcumin (Curcuma longa), Glycyrrhizin (Glycyrrhiza glabra), Eugenol (Ocimum sanctum) and Quercetin (Azadirachta indica). HPLC analysis was used to quantify marker compounds to ensure batch-to-batch consistency. TLC Fingerprinting was conducted for rapid identity and purity confirmation of all extracts.

### Formulation-Level Testing

Each formulation underwent the following tests to confirm physical, chemical, and microbiological integrity: Flowability and particle size distribution of the LUMPY-NIL herbal powder was assessed using angle of repose and sieve analysis. LUMPY-NIL herbal powder was also evaluated for uniform dispersion, taste, and odour upon mixing with water together with its pH and Disintegration to ensure optimal gastrointestinal compatibility. LUMPY-NIL dermal spray was analysed for the droplet size and dispersion pattern using a laser diffraction analyser. Accelerated stability testing was conducted at 40°C+/-2°C, with 75+/-5% RH. Microbial load and preservative efficacy was assessed to ensured sterility and prolonged shelf life. Consistency and viscosity of LUMPY-NIL ointment was assessed using a texture analyser. Accelerated stability testing was performed as per ICH guidelines (temperature and humidity control) for 6 months.

### Packaging and Labelling Standards

Packaging materials and containers (HDPE bottles for powder, glass spray bottles, laminated ointment tubes) were evaluated for compatibility, non-reactivity, and UV protection. All products were labelled with manufacturing/expiry dates, batch numbers, usage instructions, dosage, and storage conditions per regulatory requirements. These stringent quality control measures were integral to ensuring that all LUMPY-NIL formulations delivered consistent therapeutic efficacy and were safe for use in livestock.

### Study design, subjects, Clinical Examination and Data Recording

The field trial included 52 lactating cattle clinically diagnosed as LSD by the veterinarian, to evaluate the efficacy of a combination herbal therapy using three LUMPY-NIL formulations: LUMPY-NIL Herbal Powder (oral), LUMPY-NIL Dermal Spray (topical, for closed nodules), and LUMPY-NIL Ointment (topical, for open wounds). The study protocol was approved by the IAEC of Skanda Lifesciences Private Ltd and the Committee for Control and Supervision of Experiments on Animals (approval number V-11011(13)/2-CPCSEA-DADF), Government of India. Lactating cows with mild to severe clinical symptoms of LSDV, treated by veterinarians (and which the veterinarian suspected was caused by LSDV) in the practice areas of the Hoskote and Doddaballapur taluks of Bangalore Rural District and Udupi District in Karnataka state, India were included in this study. All cows included in this study were of Holstein Friesian breed, most of them being housed in the back yard of the farmer’s house. The cows were fed locally available dry fodder with additional supplementation of commercially prepared concentrate feeds. The cows were not vaccinated against LSDV and were milked twice daily.

At the time of enrolment (d 0), each cow was examined clinically by the attending veterinarian. The diagnosis of LSD infection was established based on clinical signs (fever, excessive salivation, depression, agalactia, anorexia, lacrimation, ocular and nasal discharge, emaciation due to necrotic plaques and presence of well circumscribed, slightly raised nodules, i.e., inclusion criteria). The nodules were rough, firm and found to be painful when palpated and were observed all over the body including muzzle, nasal and buccal mucosal membranes, nares, brisket region, back, legs, udder, scrotum, perineum, eyelids and lower ear. Superficial lymph nodes were enlarged in some animals with typical signs of lymphadenopathy. Lactating cow’s milk production was reduced by 50-80%. There were signs of clinical mastitis in 20 animals. During the initial visit by the veterinarian, the farmer provided informed consent by signing a consent form after the details of the study were thoroughly explained. Animals having 5-10 lumps or nodules (1 to 5 cms in diameter without any open wounds) were classified as mild case, while animals having 20-30 nodules (with few open wounds) were classified as moderate case and those with more than >30 lumps or nodules (with many nodules >5 cms and open wounds) were classified as severe case.

Pre-examination preparations included moving the affected animal to a separate clean and well-lit examination area and organisation of the necessary materials required, including the ointment, gloves, thermometer, cotton swabs, marking pens, and data recording sheets. Each animal was assigned a unique identification number and the details were recorded on the data recording sheet. The body temperature was measured using a thermometer and the examination of the skin lesions caused by lumpy skin disease on each cow was performed systematically starting from the head and progress toward the tail. The location, size, appearance, and severity of each lesion was documented on the data recording sheet. Selected lesions at the discretion of the attending veterinarian were marked using pens for future reference during follow-up examinations.

Animals were categorized into mild (n = 14), moderate (n = 20), and severe (n = 18) cases based on lesion number, size and type as mentioned above. Following examination the affected animals were treated by oral administration of LUMPY-NIL Herbal Powder at 30 g twice daily mixed with jaggery or banana, along with twice-daily topical application of the spray for closed nodules and ointment for open wounds, following cleaning of the affected areas. The dosage of oral powder was arrived based upon the IC_50_ and EC_100_ values of the selected phytochemicals. Mild cases received combination therapy for 7 days. Moderate cases received the same for 7 days, followed by oral powder once daily for an additional 7 days, with continued topical application. Severe cases followed the same initial 7-day combination treatment, followed by once-daily oral powder for 14 more days, with ongoing topical therapy until symptom resolution. The first oral dose was given by the veterinarian and the second and subsequent doses were administered by the owner following the advice by the veterinarian. All treatments were recorded by the veterinarian on the study data collection form. Clinical assessments were conducted once in 3 days with daily telephonic follow-up, monitoring for reduction in nodule size and number, cessation of new lesion formation, improvement in systemic symptoms (fever, appetite, activity), and recovery of milk yield, to evaluate therapeutic efficacy. During the follow-up visits by the veterinarian the examination process was repeated and the observations were recorded (changes in lesion appearance, size, and severity including the general clinical condition of the animal). The efficacy of the treatment was grossly visible in all treated animals. The data from all treatment records was compiled and analysed (using descriptive statistics) to quantify the efficacy and safety of the novel lumpy skin disease treatment outlined in this study.

## 3. Results

Phytochemical evaluated in this study showed binding efficacy against the selected LSDV targets at a concentration which are therapeutically feasible (Figure 1). The binding efficacy of the phytochemicals was observed against both virus entry (Table 1) and replication (Table 2) targets. The IC_50_ of the phytochemicals tested against the LSDV envelop protein and its polymerase ranged from 0.24 to 14.52 µM. The concentration response curves of the phytochemicals against the LSDV polymerase indicated their potential in effectively preventing LSDV replication (Figure 1). Based on these IC_50_ values of the phytochemicals, the different polyherbal formulations were developed to achieve optimal clinical efficacy.

**Table 1:**
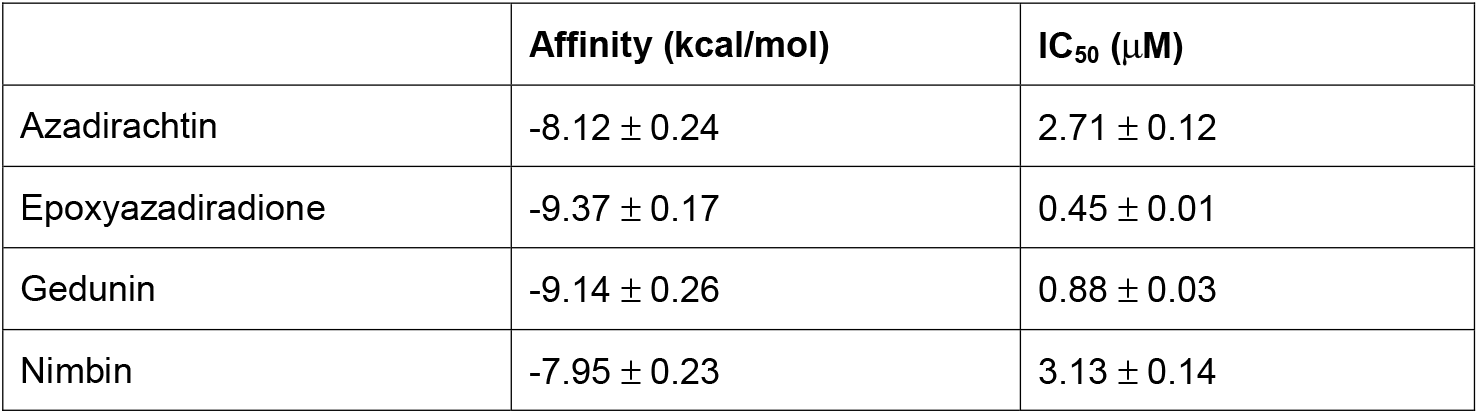
Binding affinity and IC_50_ of selected phytochemicals against LSD virus envelop protein

|  | **Affinity (kcal/mol)** | **IC<sub>50</sub> (μM)** |
| --- | --- | --- |
| Azadirachtin | -8.12 ± 0.24 | 2.71 ± 0.12 |
| Epoxyazadiradione | -9.37 ± 0.17 | 0.45 ± 0.01 |
| Gedunin | -9.14 ± 0.26 | 0.88 ± 0.03 |
| Nimbin | -7.95 ± 0.23 | 3.13 ± 0.14 |

**Table 2:**
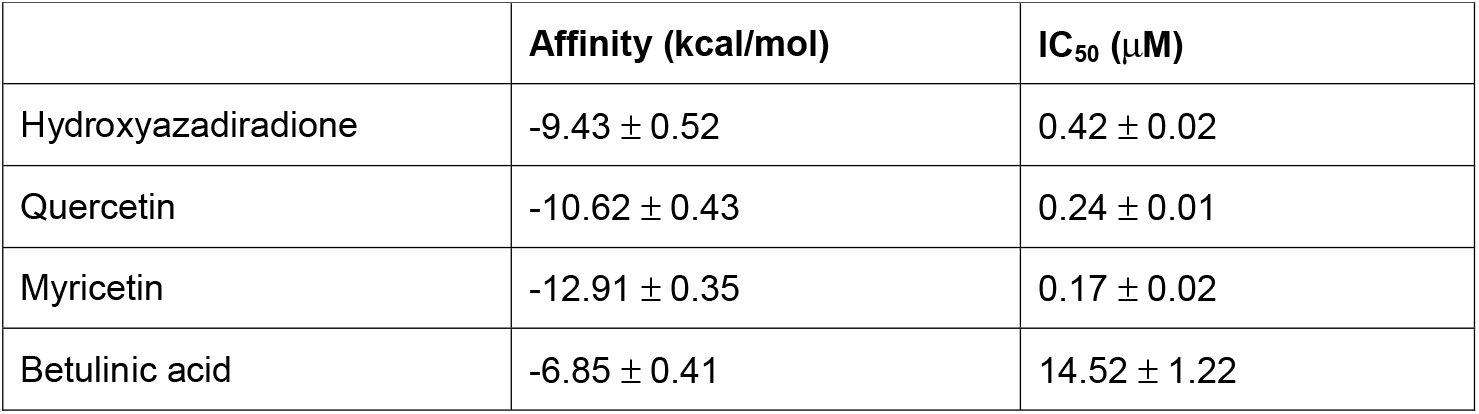
Binding affinity and IC_50_ of selected phytochemicals against LSD virus PolyAPolymerase

|  | **Affinity (kcal/mol)** | **IC<sub>50</sub> (μM)** |
| --- | --- | --- |
| Hydroxyazadiradione | -9.43 ± 0.52 | 0.42 ± 0.02 |
| Quercetin | -10.62 ± 0.43 | 0.24 ± 0.01 |
| Myricetin | -12.91 ± 0.35 | 0.17 ± 0.02 |
| Betulinic acid | -6.85 ± 0.41 | 14.52 ± 1.22 |

**Figure 1:**
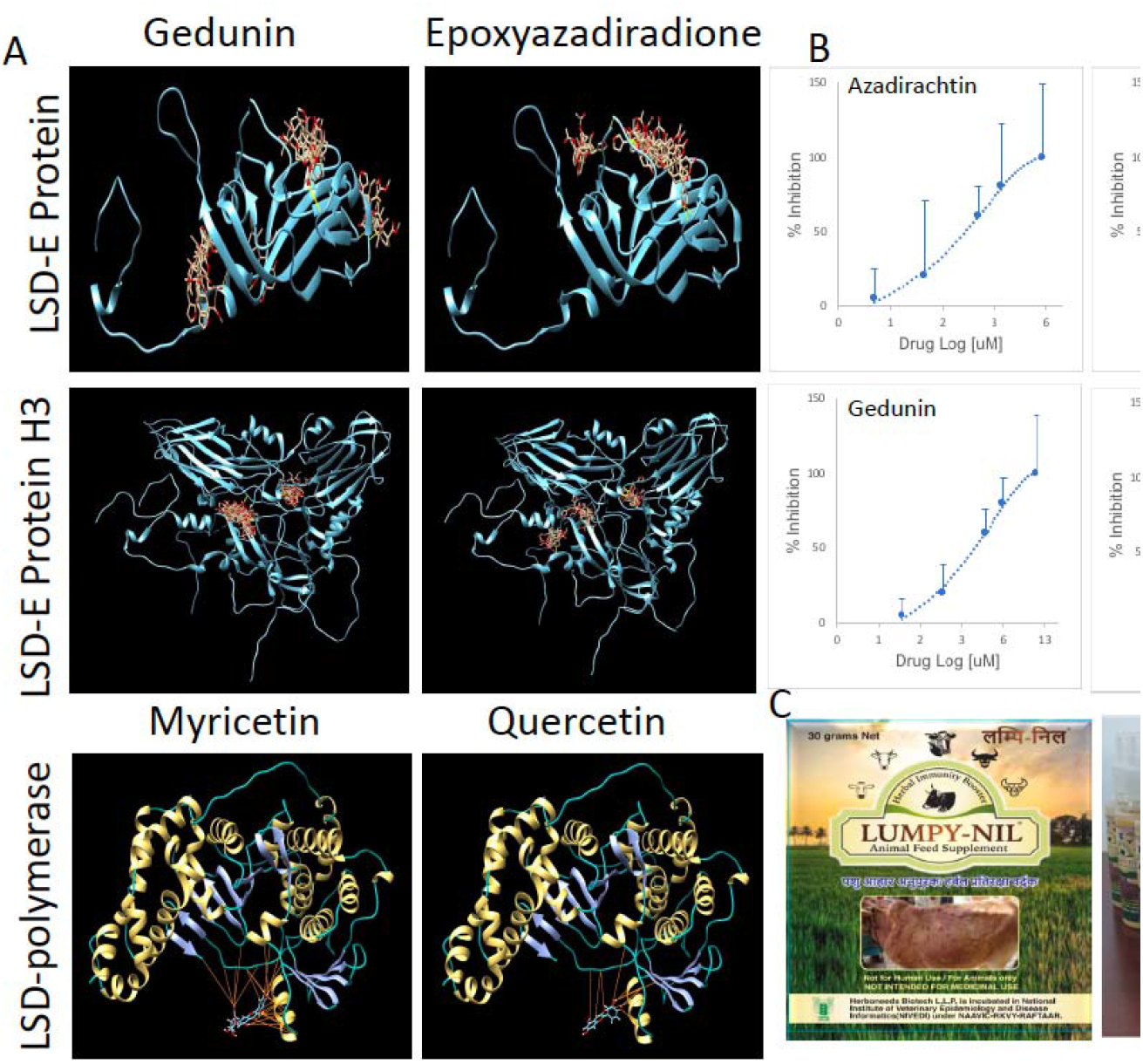
Phytochemical binding, inhibitory affinity, and formulation of herbal therapeutics against Lumpy Skin Disease Virus (LSDV). (A) Molecular docking of selected phytochemicals, Gedunin, Epoxyazadiradione, Myricetin, and Quercetin, against LSDV targets including LSDV envelope protein (LSD-E), LSD-E Protein H3, and LSDV polymerase. The ligands are shown occupying predicted binding pockets on the respective protein targets, suggesting high-affinity interactions (Yellow lines represent hydrogen bonds) that may inhibit viral entry and replication. (B) In vitro inhibition curves of four active phytochemicals (Azadirachtin, Epoxyazadiradione, Gedunin, and Nimbin) against LSDV targets. The dose-dependent response (log scale) indicates effective inhibition, with IC_50_ values ranging from 0.24 to 14.52 µM, highlighting their therapeutic feasibility. Error bars represent standard deviation (n=3). (C) Commercial formulation and product development of LUMPY-NIL, images of the packaged herbal powder and spray prepared for field application in LSD-affected cattle is shown. These formulations were designed to deliver the active phytochemicals with optimal stability and bioavailability.

### Stability assessment of the herbal formulations

A 180-day accelerated stability study was conducted for the formulated herbal ointment/spray/powder under controlled conditions (40□±□2□°C temperature and 75□±□5% relative humidity), in accordance with ICH guidelines. The formulation was evaluated at 0, 30, 60, 90, and 180 days for its physical, chemical, and microbiological parameters. Physical characteristics, including colour and texture, remained consistent throughout the study period. The ointment retained its greenish-brown colour and soft semi-solid consistency across all time points, indicating no phase separation, drying, or physical degradation. Organoleptic properties of the herbal spray, including appearance and consistency, remained stable throughout the test period. The formulation consistently appeared as a greenish-yellow, viscous liquid with visible separation of oil particles across all time points, indicating no significant phase instability under accelerated storage conditions. Moisture content of the ointment showed minor variation, with an initial value of 2.98% on day 0, reducing to 2.33% at day 30, and then gradually increasing to 2.81% by day 180. These fluctuations remained within acceptable pharmaceutical limits, indicating adequate moisture stability. Moisture content of the spray showed initial decline from 45.46% at day 0 to 34.76% by day 30, after which it remained relatively stable, ranging between 35.22% and 35.69% through day 180. This early drop may indicate equilibration under test conditions, with subsequent stabilization reflecting adequate formulation robustness. pH values remained stable, fluctuating within a narrow range from 5.326 on day 0 to 5.492 on day 180. This consistency indicates good chemical stability and suggests the formulation remains within the ideal pH range for topical application, minimizing the risk of skin irritation. Viscosity increased progressively from 25.28 Pa·s on day 0 to 45.5 Pa·s by day 180. The formulation remained homogenous and easily applicable throughout the study period despite this thickening trend, suggesting no adverse impact on usability. Rancidity was noted as slightly oxidized across all intervals, with no observed increase in oxidative degradation over time. This indicates stable lipid components in the formulation. Chemical stability was confirmed through High Performance Thin Layer Chromatography (HPTLC) fingerprinting, which showed compliance at all testing intervals. The consistent phytochemical profile suggests that the active constituents of the herbal extracts remained stable over the course of the study. Microbiological analysis revealed a total plate count of 130 cfu/g and yeast mould count of 210 cfu/g on day 0. These values markedly reduced to <10 cfu/g from day 30 onward and remained consistently low throughout the stability assessment period. This improvement in microbial load suggests the effectiveness of preservatives and good formulation stability under the accelerated conditions. Based on the cumulative data, all the three formulations demonstrated robust stability under accelerated conditions for 180 days. The results indicated a projected shelf life of up to 24 months under recommended storage conditions.

### Efficacy of herbal formulations in field trials

The clinical cases were treated with all three formulations for optimal outcome. Following the initiation of the treatment none of the cutaneous nodules were observed to be bleeding and cessation of new nodules appearance was observed within 2-4 days (Figures 2 and 3). Symptoms of fever, restlessness and loss of appetite improved immediately and was restored to normal within 3-4 days following treatment initiation. Loss of milk yield was the slowest to recover, with 20% recovery in milk yield observed by day 7 of treatment with complete recovery of milk yield observed by 15^th^ day of treatment (Figure 2). A progressive improvement in milk yield was observed across all clinical categories (mild, moderate, and severe) following initiation of treatment. The heatmap (Figure 2) illustrates a clear trend of recovery, beginning around day 6–7 in mildly affected animals, with most returning to baseline milk production by day 13–14. Moderately affected animals showed gradual recovery, with milk yield normalization occurring between days 14 and 17. In severely affected cases, recovery began slightly later but followed a consistent upward trend, with full recovery achieved in most animals by day 17–21 (Figure 2).

**Figure 2:**
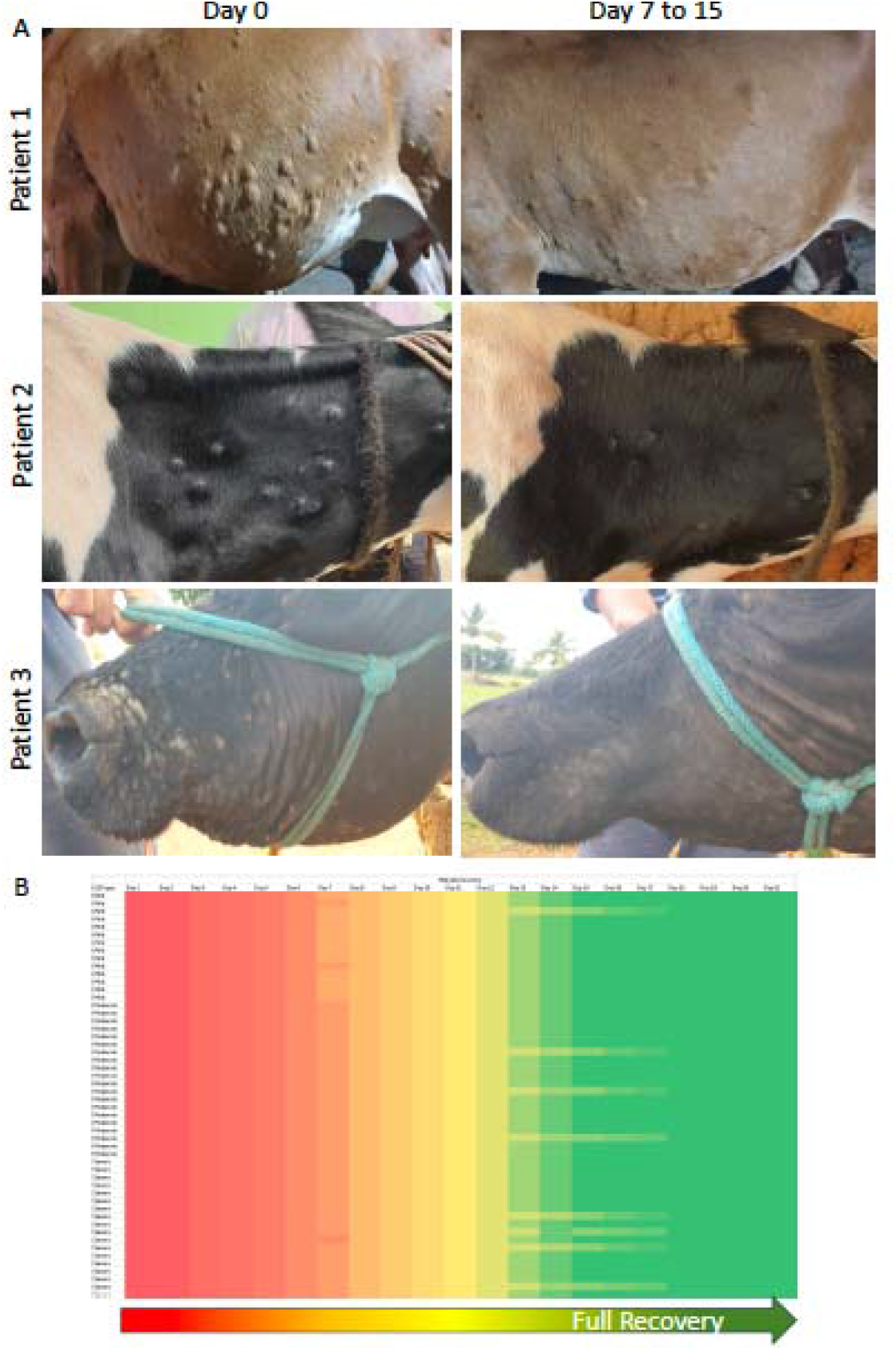
Clinical recovery of LSD-affected cattle following treatment with polyherbal formulations. (A) Representative images of three LSDV-infected cattle (Patients 1–3) showing visible skin lesion regression following treatment in various parts of the body. Images compare the severity of cutaneous nodules at Day 0 (pre-treatment) and post-treatment days 7 to 15, demonstrating substantial healing, drying, and resolution of nodules. Complete or near-complete recovery is evident by Day 15. (B) Heatmap showing temporal trends in milk yield recovery among treated animals across clinical severity categories (mild, moderate, severe). Each row represents an individual animal, and colour intensity corresponds to the degree of recovery, progressing from red (low yield) to green (full recovery). A consistent improvement in milk production is observed, with mildly affected animals recovering by Day 13-14 and severely affected cases showing complete recovery by Day 15-18.

**Figure 3:**
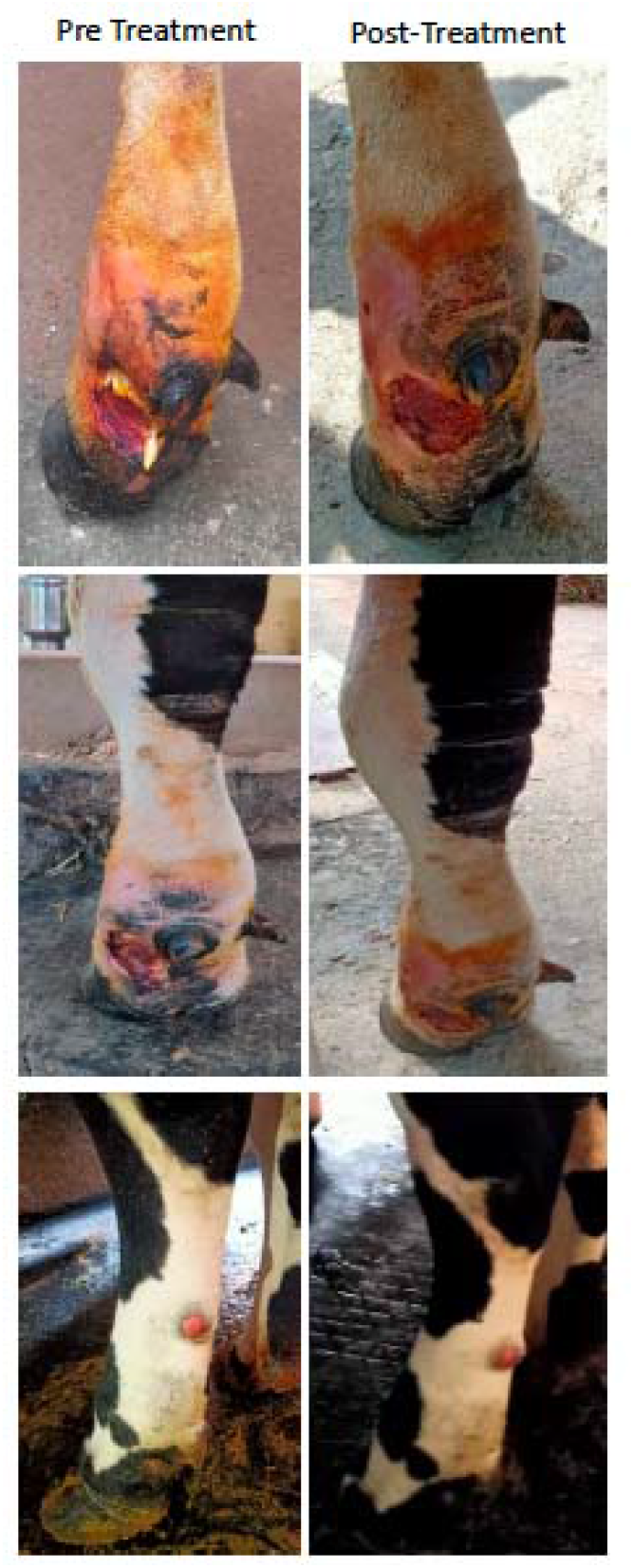
Clinical recovery of cutaneious wounds following treatment with LUMPY-NIL ointment/spray. Representative images of three LSDV-infected cattle with cutaneious wounds pre and post treatment with LUMPY-NIL ointment/spray. Images on Day 0 (left panels) depict prominent ulcerative cutaneous lesions. By Days 7 to 15 (right panels), notable improvement was observed following daily topical application of LUMPY-NIL ointment/spray. Many of the wounds had entered the re-epithelialization phase, characterized by visible contraction of wound margins and the emergence of new epidermal tissue, indicating the initiation of tissue repair and regeneration.

The cutaneous nodules exhibited clear signs of regression beginning as early as day 4 of treatment, with noticeable drying and shrinkage evident in the majority of animals by day 7. Over the course of therapy, the nodules progressively desiccated, sloughed off, and underwent epithelial healing without secondary complications. By day 10, a marked reduction in both the size and number of nodules was observed across all cases, with an efficacy rate ranging between 81% and 94% in terms of nodule resolution recorded between days 10 and 15 of treatment. Importantly, no new nodules were observed to develop during the treatment period, indicating strong therapeutic efficacy and containment of viral progression. By the 15th day, nearly all treated animals displayed complete resolution of visible nodules, with the skin returning to normal or near-normal appearance (Figure 2). These findings suggest that the polyherbal treatment not only promotes healing of existing lesions but also effectively halts the emergence of new ones. Remarkable healing of open wounds was observed in all treated animals following the initiation of the combination therapy with LUMPY-NIL herbal powder, dermal Spray, and ointment. Initial signs of wound recovery, such as a noticeable reduction in local inflammation, drying of the wound bed, and the appearance of healthy granulation tissue were evident as early as day 4 of treatment. The healing process progressed consistently, with granulation tissue gradually maturing and transitioning into well-organized epithelial layers. By day 10, many wounds had entered the re-epithelialization phase, characterized by visible contraction of wound margins and the emergence of new epidermal tissue (Figure 3). Complete healing, defined by full epithelial closure and restoration of skin integrity without scabbing or residual lesions, was observed in the majority of animals by day 14. Notably, no cases of secondary bacterial infection, exudation, or delayed healing were reported throughout the course of treatment. These findings validate the effectiveness, safety, and tolerability of the topical applications in accelerating wound resolution and promoting robust tissue regeneration in LSD-affected bovines.

## 4. Discussion

Lumpy Skin Disease (LSD), caused by the Capripoxvirus LSDV, continues to challenge cattle health management worldwide, especially in endemic regions and newly affected zones. Traditional control measures have largely centred around vaccination and symptomatic treatment.[3,6] Although preventive vaccination remains the cornerstone of LSD control, its effectiveness is often variable, especially during outbreaks, and there are no antiviral therapies currently approved for treatment once clinical disease manifests. In this context, the present study introduces a novel, phytochemical-based therapeutic strategy that offers both antiviral and lesion-resolving properties, representing a potential paradigm shift in LSD management. Current live attenuated vaccines, such as those derived from the Neethling strain or heterologous sheep pox or goat pox viruses, have demonstrated efficacy in reducing outbreak severity and disease transmission.[2,15,16] However, these vaccines are not without limitations. Breakthrough infections have been reported even in vaccinated populations, likely due to antigenic variability, improper cold chain maintenance, or incomplete immune coverage.[13,34,35] Moreover, live vaccines can induce post-vaccinal reactions, including local swelling, fever, and even the development of LSD-like skin lesions, complicating differential diagnosis during outbreaks.[36] Additionally, vaccination does not offer therapeutic benefit once clinical symptoms appear, leaving a significant gap in disease management.

The current study addresses this therapeutic void by evaluating the antiviral efficacy of select phytochemicals that demonstrated strong binding affinity to LSDV envelope and replication proteins at concentrations ranging from 0.24 to 14.52 µM. These IC_50_ values are therapeutically achievable and suggest dual activity against viral entry and cytoplasmic replication. Compared to existing literature, where few compounds have been experimentally validated against LSDV targets, this represents an innovative attempt to identify naturally derived molecules that interfere with both the attachment to glycosaminoglycans and DNA polymerase-driven replication, which are critical steps in the viral life cycle. The clinical findings from field application further underscore the therapeutic potential of these compounds when formulated into a combination of ointment, spray, and powder. Within 2–4 days of initiating treatment, no new nodules were observed, contrasting sharply with the natural progression of LSD, where new lesions typically appear for up to 10 days post-onset.[9,19,36] Cutaneous nodules in treated animals showed rapid drying and sloughing beginning on day 4, with an 81–94% reduction in lesion burden between days 10 and 15. This rate of recovery is significantly faster than the 3–6 weeks of lesion persistence typically described in untreated cases or those managed solely with NSAIDs and antibiotics.[9,37]

In addition to cutaneous recovery, systemic symptoms such as fever, anorexia, and lethargy resolved within 3–4 days post-treatment, indicating potential early antiviral or immunomodulatory effects. To the best of our knowledge such rapid recovery has not been reported by any of the current approaches used for the clinical management of LSD. Furthermore, milk yield recovery, often delayed or incomplete in conventionally managed outbreaks,[6,38] was seen to begin as early as day 6 in mildly affected animals, with complete restoration in most cases by day 15–21. These functional outcomes are rarely reported in the context of vaccine-based management and highlight the broader impact of therapeutic intervention on productivity. Secondary bacterial infections of skin lesions are a common complication in LSD, often leading to delayed wound healing, prolonged morbidity, and increased antibiotic usage.[19,37] In this study, the application of the polyherbal spray and ointment formulations was associated with excellent wound healing outcomes, with no signs of secondary infection, exudation, or delayed recovery observed throughout the treatment period. The rapid resolution of inflammation, promotion of granulation tissue, and consistent epithelialization suggest strong antimicrobial and anti-inflammatory properties of the topical LUMPY-NIL formulations. These findings indicate that the herbal spray and ointment not only accelerated lesion healing but also effectively controlled secondary infections, reducing the need for systemic antibiotics and supporting safer, more sustainable therapeutic practices in LSD management.

The therapeutic efficacy of LUMPY-NIL observed in cattle raises the possibility that this phytochemical-based formulation may be effective across other susceptible species, such as giraffes and impalas, which are also known to be affected by LSDV.[39,40] Given the conserved structure and replication strategy of LSDV, an enveloped virus with brick-shaped geometry and a linear dsDNA genome of approximately 154 kb, LUMPY-NIL may exert similar antiviral effects across species. The virus initiates infection by binding to host glycosaminoglycans, followed by endocytosis and a cytoplasmic replication cycle involving early, intermediate, and late gene expression phases. Phytochemicals in LUMPY-NIL, which demonstrated high binding affinity to both the viral envelope proteins and the viral DNA polymerase, could potentially interfere with these conserved mechanisms, disrupting viral attachment and inhibiting replication. Since these key viral processes are not species-specific, it is plausible that the same molecular targets are accessible in a range of host species. Hence, LUMPY-NIL may offer a cross-species therapeutic benefit, although this hypothesis warrants further investigation through comparative pharmacological studies and clinical trials in non-bovine hosts.

The phytochemical formulations also exhibited excellent pharmaceutical stability. A 180-day accelerated stability study demonstrated consistent physical, chemical, and microbiological parameters across all three formulations. HPTLC profiling confirmed the retention of active constituents, while pH and viscosity remained within acceptable ranges for dermal application. These findings suggest that the formulations are not only efficacious but also scalable and suitable for deployment in diverse field conditions, including regions with limited veterinary infrastructure. Nonetheless, certain limitations must be acknowledged. While phytochemical-based formulations provide therapeutic benefit during active disease, they do not offer long-term immunity. In contrast, vaccines, despite their limitations remain the most effective tool for population-level disease prevention and herd immunity. Moreover, this study did not evaluate the potential prophylactic effects of these phytochemicals, nor did it assess their compatibility with existing vaccination programs. Another important consideration is regulatory standardization and quality control, which will be critical for the widespread adoption of herbal therapeutics in veterinary practice.

## 5. Conclusions

In conclusion, this study highlights the promising role of phytochemical-based therapy as a complementary tool in LSD management. By offering both antiviral and wound-healing properties, these formulations can fill the therapeutic void left by vaccines and symptomatic care. The integration of such natural compounds into LSD management protocols, potentially alongside vaccination may offer a more comprehensive and sustainable approach to controlling this economically significant disease. Future studies should focus on dose standardization, long-term safety, and synergistic evaluation with vaccines to fully realize the potential of phytochemical therapeutics in trans-boundary animal disease control.

## Author Contributions

For research articles with several authors, a short paragraph specifying their individual contributions must be provided. The following statements should be used “Conceptualization, AHSK and BMR.; methodology, AHSK, VS and BMR.; software, AHSK; validation, AHSK, and VS.; formal analysis, AHSK.; Field trials, BMR.; resources, AHSK, VS and BMR.; data curation, AHSK and BMR.; writing—original draft preparation, AHSK.; writing—review and editing, AHSK, VS.; visualization, X.X.; supervision, BMR.; project administration, AHSK, and VS.; funding acquisition, VS. All authors have read and agreed to the published version of the manuscript.

## Funding

This research received no external funding

### Institutional Review Board Statement

The study protocol was approved by the IAEC of Skanda Lifesciences Private Ltd and the Committee for Control and Supervision of Experiments on Animals (approval number V-11011(13)/2-CPCSEA-DADF), Government of India.

### Informed Consent Statement

Informed consent was obtained from all patient owners prior to start of this study.

## Abbreviations

The following abbreviations are used in this manuscript:

LSD: Lumpy Skin Disease
LSDV: Lumpy skin disease virus
PCR: polymerase chain reaction
ELISA: enzyme-linked immunosorbent assay
API: Active phytochemical ingredients

